# Predicting the future development of mild cognitive impairment in the cognitively healthy elderly

**DOI:** 10.1101/2020.07.30.227496

**Authors:** Bryan A. Strange, Linda Zhang, Alba Sierra-Marcos, Eva Alfayate, Jussi Tohka, Miguel Medina

## Abstract

Identifying measures that predict future cognitive impairment in healthy individuals is necessary to inform treatment strategies for candidate dementia-preventative and modifying interventions. Here, we derive such measures by studying “converters” who transitioned from cognitively normal at baseline to mild-cognitive impairment (MCI) in a longitudinal study of 1213 elderly participants. We first establish reduced grey matter density (GMD) in left entorhinal cortex (EC) as a biomarker for impending cognitive decline in healthy individuals, employing a matched sampling control for several dementia risk-factors, thereby mitigating the potential effects of bias on our statistical tests. Next, we determine the predictive performance of baseline demographic, genetic, neuropsychological and MRI measures by entering these variables into an elastic net-regularized classifier. Our trained statistical model classified converters and controls with validation Area-Under-the-Curve>0.9, identifying only delayed verbal memory and left EC GMD as relevant predictors for classification. This performance was maintained on test classification of out-of-sample converters and controls. Our results suggest a parsimonious but powerful predictive model for MCI development in the cognitively healthy elderly.

---

Alzheimer’s disease (AD) is the most common form of dementia and is currently estimated to affect more than 46 million people worldwide, with prevalence predicted to rise to over 130 million by 2050^1^. It has been established that certain pathophysiological hallmarks of AD (neurofibrillary tangles and amyloid plaques) emerge decades before the first manifestations of clinically observable dementia^2-4^, indicating that biomarkers of AD are likely present in those individuals that will develop AD even when they are cognitively normal. Thus, the clinical disease stages of AD can be divided into three phases^4^: 1) a presymptomatic phase, in which individuals are cognitively normal but already exhibit AD pathological changes, 2) a prodromal phase of AD, which overlaps with mild cognitive impairment (MCI), characterized by early cognitive symptoms (typically deficits in episodic memory) not severe enough to meet the criteria for dementia, 3) the dementia phase, in which multiple domains of cognition are impaired to the extent that the patient experiences loss of daily function. While currently there are limited treatment options available in AD, it is likely that future strategies will be most effective if applied at the earliest stage of the disease. Consequently, identification of well-characterized measures that manifest early and can track the AD process are necessary to inform treatment strategies for candidate preventative and disease-modifying interventions^5-7^. Indeed, the on-going A4 study^8^ aims to identify cognitively normal individuals with amyloid accumulation and treat with anti-amyloid therapy.

Certain AD risk factors, such as increasing age, fewer years of education and the apolipoprotein E (APOE) ε4 allele, are well recognised^9^. However, the relative contribution of each of these factors to the likelihood of development of AD during a particular time-period, and from a defined cognitive starting point (*e*.*g*., within normal limits on standard neuropsychological testing), is currently unknown. Large-scale studies are required to identify additional predictive indices that can be determined non-invasively and relatively routinely, such as neuropsychological tests or structural magnetic resonance imaging (MRI). The majority of large-scale longitudinal studies have focused on predicting the transition of MCI to AD^10-12^. By contrast, studies examining healthy to MCI transition are currently limited^13-19^, although relevant information has been obtained from presymptomatic studies of autosomal dominant familial AD studies^20-23^. The challenge is to determine which parameters show most discrimination between cognitively normal individuals destined for MCI and eventually to sporadic AD *vs*. those that remain healthy.

A further difficulty relates to the causal chain of the AD process whereby known risk factors, such as age and APOEε4 genotype, in turn may influence neuropsychological performance and brain structure^24-26^. On the one hand, in assessing the efficacy of brain imaging as a predictor of future MCI conversion, we must appropriately sample or otherwise net out the confounding contribution of these risk factors to the prediction problem. On the other, we might wish to recruit all available subject attributes for the purposes of improving prediction. The latter scenario requires a statistical framework that can account for the likely correlation between subject attributes (*e*.*g*., medial temporal lobe structural integrity is known to correlate with memory test scores in the elderly^27^) and that can select those variables that maximize predictive power while pruning those that either contain no discriminant power or that are redundant in relation to their predictive contribution.

To address the first of these issues, we employed techniques from matched sampling^28^. Matching is a non-parametric pre-processing method that reduces covariate imbalance between groups rendering the treatment and outcome variables independent (or almost independent) of one another. Matching has several useful implications: improved causal inference, heightened power, and a reduced sensitivity to model specification. This strategy is typically applied in the comparison of treatment and control groups, and in case-control cohort designs, such as in biomarker research studies where controls can be selected that match cases on risk factors for the outcome. To address the second issue, accounting for likely correlation between subject attributes, we developed a classification model based on the Elastic net^29^. Elastic nets are advantageous for classification where numbers of predictors are large relative to the number of subjects. Elastic net optimization combines classification with an implicit feature selection step, tending toward retaining small numbers of isolated predictors while at the same time preserving groups of correlated features, if such structure exists between them.

We applied these techniques to data from the Vallecas Project^30^, a single-site, community-based, longitudinal study on a recruited pool of 1,213 individuals aged 69-86 and followed up at yearly intervals. At each visit, volunteers undergo detailed neuropsychological and clinical evaluation, and 3 Tesla (3T) MRI, to assess non-invasively the macroscopic consequence of neuronal damage occurring in neurodegeneration. In the first analysis approach, we determined, using standard difference of means estimates, which of the demographic, genetic, neuropsychological and MRI variables significantly distinguished between the two groups of cognitively healthy individuals, with group membership defined by whether they developed MCI the following year or remained cognitively normal. Since our dataset is observational in its native form, it is not balanced in any of the measured background covariates. We therefore apply matching techniques to identify a well-matched control group through the selection of subjects from a control reservoir using a large spectrum of relevant background variables to match upon. Our second approach invokes the elastic net classifier^29^ on a large number of clinical, demographic and imaging variables to automatically assess individuals according to their probability of future conversion to MCI. Both approaches were applied to Vallecas project participants who converted from cognitively normal to MCI from Visit 1 to 2. We subsequently applied the classifier to a test sample who converted to MCI in later visits.

## RESULTS

### Cognitively healthy individuals destined for MCI

From a pool of 1213 participants, specific criteria were applied to select the cases for the present study (**Figure 1**). To approximate cognitive normality at baseline (visit 1; V1) we selected participants with Clinical Dementia Rating (CDR)=0 and mini-mental state examination (MMSE)>26. We first focused on those cognitively healthy individuals developing MCI from V1 to V2. Participants who were considered converters in V2, but returned to a healthy state in V3 (*i*.*e*., ‘reverters’ in V3) were excluded, leaving 813 participants (63.25% females). By evaluating diagnostic status at V2 and V3, 23 participants were considered future converters (evolving from a cognitive normal state in V1 to a state of MCI in V2, which persisted at V3), and 790 were considered controls (non-converters during this two year period). The number of converters is in line with incident rates in other populations^3,31-32^. Of the 23 converters, 11 developed amnestic MCI and 12 multi-domain MCI. For each converter the closest match in terms of APOE genotype, gender, age, years of education, MMSE and total intracranial volume (TIV; determined from structural MRI) was identified, creating a subgroup of 23 matched controls.

**Figure 1.**
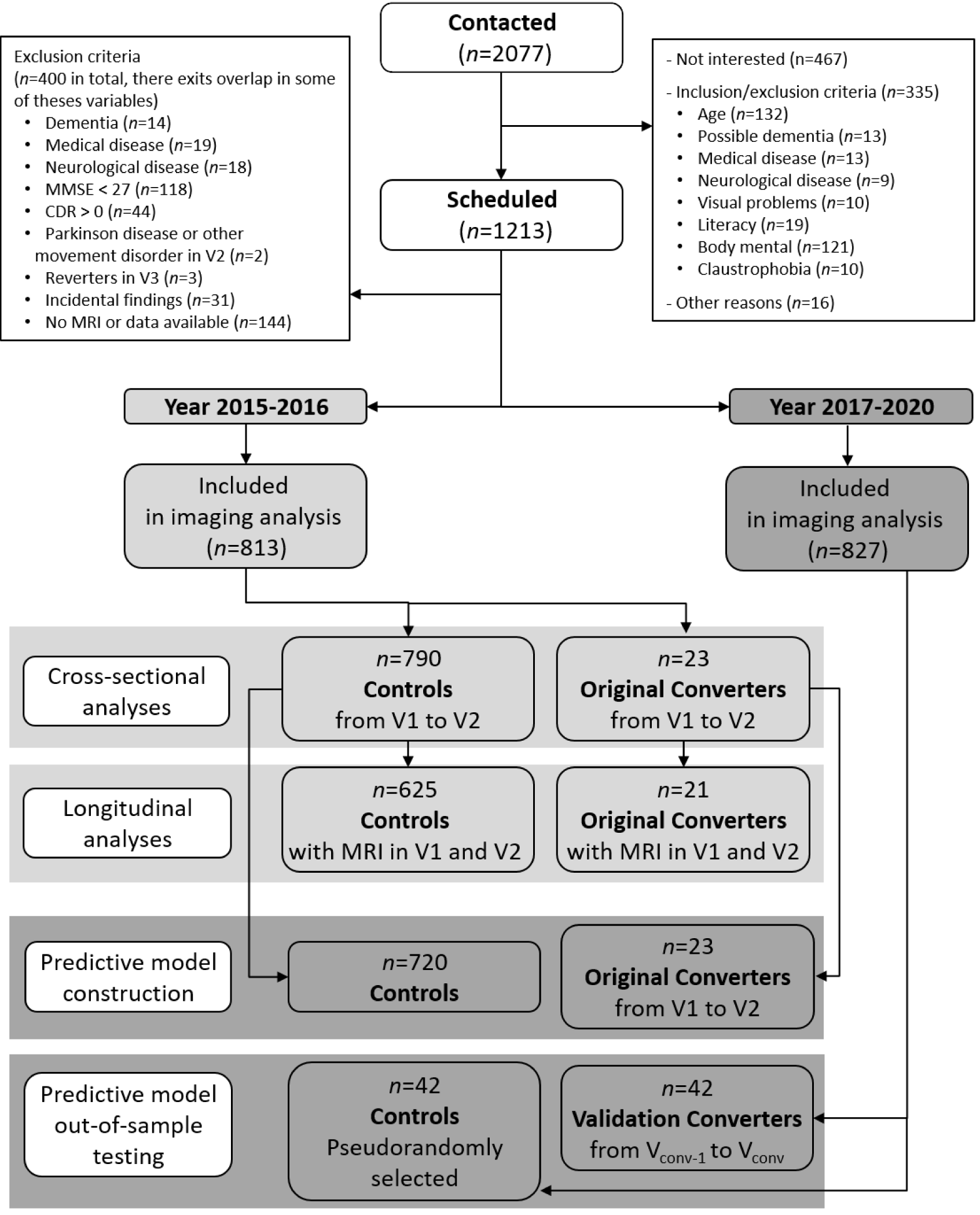
Flowchart of participant recruitment from the Vallecas Project. The exclusion criteria applied to the current study and ensuing groups of converters and controls are indicated. Note that 827 participants were included in the 2017-2020 analyses, as converters from V2 to V3, or controls not yet attending V3, were excluded from the control group in our 2015-2016 analyses.

### Biomarkers associated with impending cognitive decline

APOEε4 load was higher in converters *vs*. controls (**Table 1**), in line with the known risk conferred by this allele in developing AD^9^. There was also a significant effect of gender, reflecting more male converters in a predominantly female study population. Although converters and controls were clinically indistinguishable at V1, and all their cognitive scores were above the 20^th^ centile corrected for age (69-71 years)^33^, V1 scores for subsequent converters *vs*. all controls were significantly reduced on delayed verbal memory testing (Free and Cued Selective Reminding Test, FCSRT, delayed total recall). By contrast, delayed non-verbal memory (Rey–Osterrieth Complex Figure) scores did not differ between groups. Critically, the FCSRT and functional activities questionnaire (FAQ) test scores remained significantly different following comparison of the 23 converters to the 23 matched controls (**Table 1**).

**Table 1.**
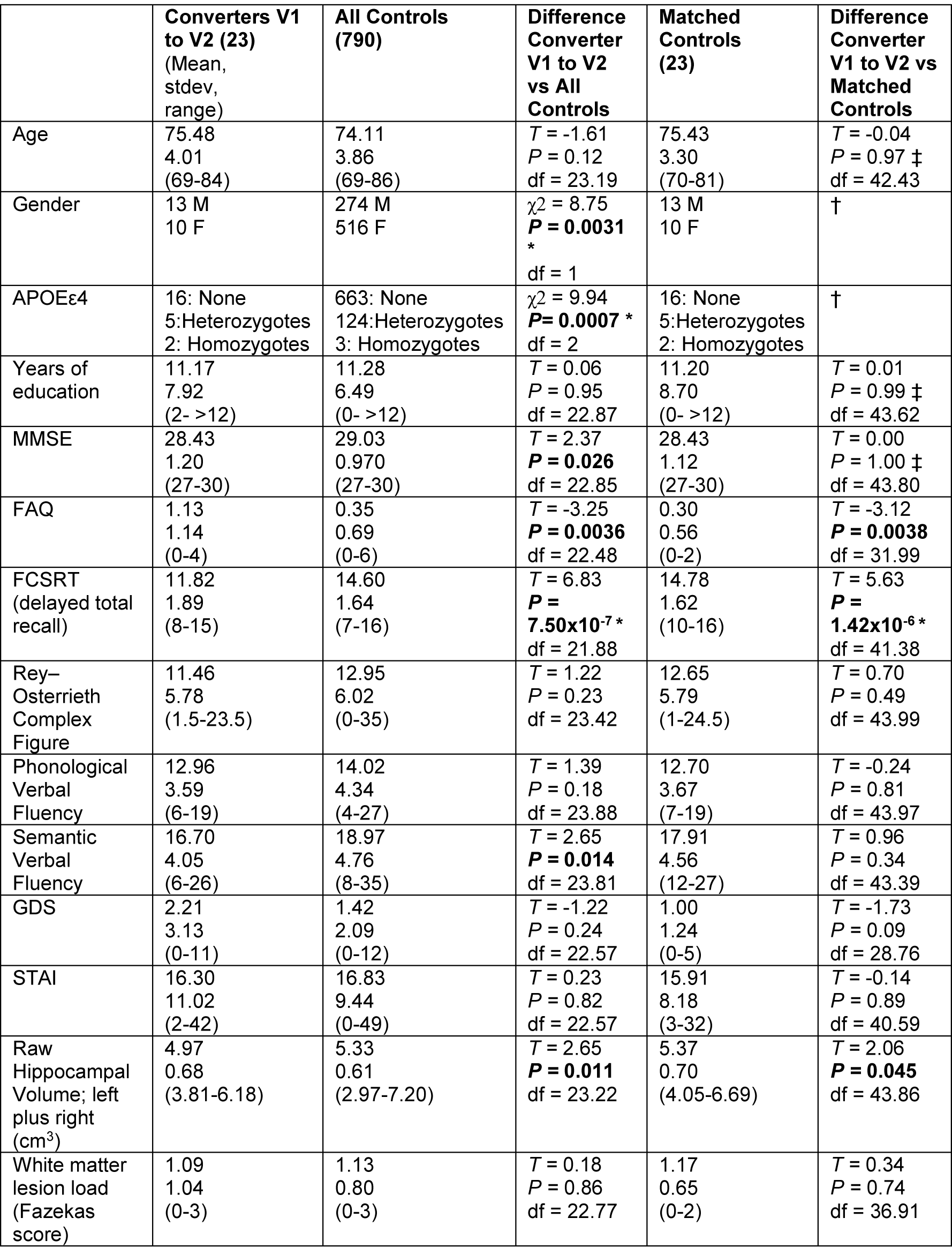
Demographic, genetic and neuropsychological participant values, as well as hippocampal volumes and Fazekas white matter integrity score, at Visit 1. Abbreviations: APOEε4: APOEε4 allele, df: degrees of freedom, FAQ: Functional Activities Questionnaire, FCSRT: Free and Cued Selective Reminding Test, GDS: Geriatric Depression Scale, M/F: Males/Females, MMSE: Mini Mental State Examination, STAI: State-Trait Anxiety Inventory - Trait, stdev: standard deviation. †These values are obtained from exact matched-samples, so by definition are not different. ‡Values obtained after propensity score matching, which included this variable. Significant *P*-values are given in bold. *T* values pertain to Welch’s *t*-test (two-tailed). *Survives Bonferroni correction for the 14 tests. There was no correlation between hippocampal volume and total intracranial volume across converters and controls (Pearson’s r = 0.043; *P* = 0.217), so raw hippocampal volumes are presented.

Hippocampal volume was not strongly modulated by future MCI development (**Table 1**), with the observed difference not surviving correction for multiple comparisons either when comparing against all controls or the matched control subgroup. White matter lesion load, indexed by the Fazekas score^34^, showed no difference between future converters and controls. By contrast, whole-brain voxel-wise analysis of grey matter density (GMD) showed reduced GMD in converters *vs*. all controls selectively in the medial temporal lobe (**Figure 2a-c; Supplementary Table 1**). Effects were observed in bilateral amygdala, bilateral anterior hippocampus, and left entorhinal cortex (EC). A significant cluster in EC was observed, with a second cluster extending more anteriorly in, or near, transentorhinal cortex. Critically, in the comparison between converters and matched controls (**Figure 2d-e**), the only brain region surviving whole-brain family-wise error (FWE) correction at *P*<0.05 was within EC (**Supplementary Table 2**). There was no significant difference in EC GMD between the 11 converters subsequently developing amnestic MCI (mean; s.e.m. = 0.53; 0.01) and 12 multi-domain MCI (0.55; 0.01) (*t*(21) = −1.319, *P* = 0.201).

**Table 2.** Demographic, genetic and neuropsychological participant values, as well as hippocampal volumes and Fazekas white matter integrity score, of test converters. Note that the reduction in size of the control group (now 762) reflects the fact that some test converters were originally controls at V1 (their V1 data have been excluded to preclude mixing between- and within-subject comparisons). Abbreviations are as for Table 1. Significant *P*-values are given in bold. *T* values pertain to Welch’s *t*-test (two-tailed). *Survives Bonferroni correction for the 14 tests.

|  | **Test Converters (42)**<br>(Mean,<br>stdev,<br>range) | **All Controls**<br>**(762)** | **Difference Test**<br>**Converters vs.**<br>**All Controls** | **Difference Original**<br>**V1-V2 Converters**<br>**vs. Test Converters** |
| --- | --- | --- | --- | --- |
| Age | 77.43<br>4.31<br>(71-87) | 74.09<br>3.84<br>(69-86) | $T = -4.91$<br>$P =$<br>**$1.0 \times 10^{-5} *$**<br>$df = 44.67$ | $T = 1.83$<br>$P = 0.07$<br>$df = 48.24$ |
| Gender | 14 M<br>28 F | 264 M<br>498 F | $\chi^2 = 0.03$<br>$P = 0.86$<br>$df = 1$ | $\chi^2 = 3.29$<br>$P = 0.07$<br>$df = 1$ |
| APOEε4 | 27: None<br>14: Heterozygotes<br>1: Homozygote | 643: None<br>116: Heterozygotes<br>3: Homozygotes | $\chi^2 = 13.16$<br>**$P = 0.001 *$**<br>$df = 2$ | $\chi^2 = 1.54$<br>$P = 0.46$<br>$df = 2$ |
| Years of education | 10.74<br>5.66<br>(0-24) | 11.23<br>6.49<br>(0-50) | $T = 0.55$<br>$P = 0.59$<br>$df = 47.14$ | $T = -0.23$<br>$P = 0.82$<br>$df = 34.55$ |
| MMSE | 28.40<br>1.06<br>(27-30) | 29.04<br>0.969<br>(27-30) | $T = 3.78$<br>**$P = 0.0005 *$**<br>$df = 44.85$ | $T = -0.10$<br>$P = 0.92$<br>$df = 40.85$ |
| FAQ | 1.21<br>1.14<br>(0-4) | 0.34<br>0.68<br>(0-6) | $T = -4.94$<br>**$P = 0.0001 *$**<br>$df = 42.66$ | $T = 0.28$<br>$P = 0.78$<br>$df = 45.30$ |
| FCSRT | 13.38<br>2.17<br>(9-16) | 14.64<br>1.61<br>(7-16) | $T = 3.70$<br>**$P = 0.0006 *$**<br>$df = 43.52$ | $T = 2.98$<br>**$P = 0.005$**<br>$df = 48.25$ |
| Rey-Osterrieth Complex Figure | 10.67<br>7.73<br>(0-36) | 13.04<br>6.01<br>(0-35) | $T = 1.46$<br>$P = 0.16$<br>$df = 22.82$ | $T = -0.39$<br>$P = 0.70$<br>$df = 40.73$ |
| Phonological Verbal Fluency | 13.90<br>4.94<br>(6-24) | 14.01<br>4.34<br>(4-27) | $T = 0.13$<br>$P = 0.90$<br>$df = 44.57$ | $T = 0.89$<br>$P = 0.38$<br>$df = 57.90$ |
| Semantic Verbal Fluency | 15.93<br>33.78<br>(9-26) | 19.05<br>4.74<br>(8-35) | $T = 5.14$<br>**$P = 5.0 \times 10^{-6} *$**<br>$df = 48.40$ | $T = -0.75$<br>$P = 0.46$<br>$df = 42.74$ |
| GDS | 2.43<br>2.57<br>(0-9) | 1.38<br>2.04<br>(0-12) | $T = -2.59$<br>**$P = 0.013$**<br>$df = 43.91$ | $T = 0.28$<br>$P = 0.78$<br>$df = 38.38$ |
| STAI | 14.78<br>10.46<br>(1-39) | 16.78<br>9.41<br>(0-49) | $T = 1.20$<br>$P = 0.24$<br>$df = 43.58$ | $T = -0.54$<br>$P = 0.59$<br>$df = 43.69$ |
| Raw Hippocampal Volume - left plus right (cm<sup>3</sup>) | 4.99<br>0.62<br>(4.07-6.53) | 5.34<br>0.61<br>(2.97-7.20) | $T = 3.56$<br>**$P = 0.0009 *$**<br>$df = 45.41$ | $T = 0.12$<br>$P = 0.91$<br>$df = 42.22$ |
| Fazekas score for white matter lesions | 1.38<br>0.96<br>(0-3) | 1.11<br>0.79<br>(0-3) | $T = -1.78$<br>$P = 0.08$<br>$df = 44.11$ | $T = 1.12$<br>$P = 0.27$<br>$df = 42.40$ |

**Figure 2.**
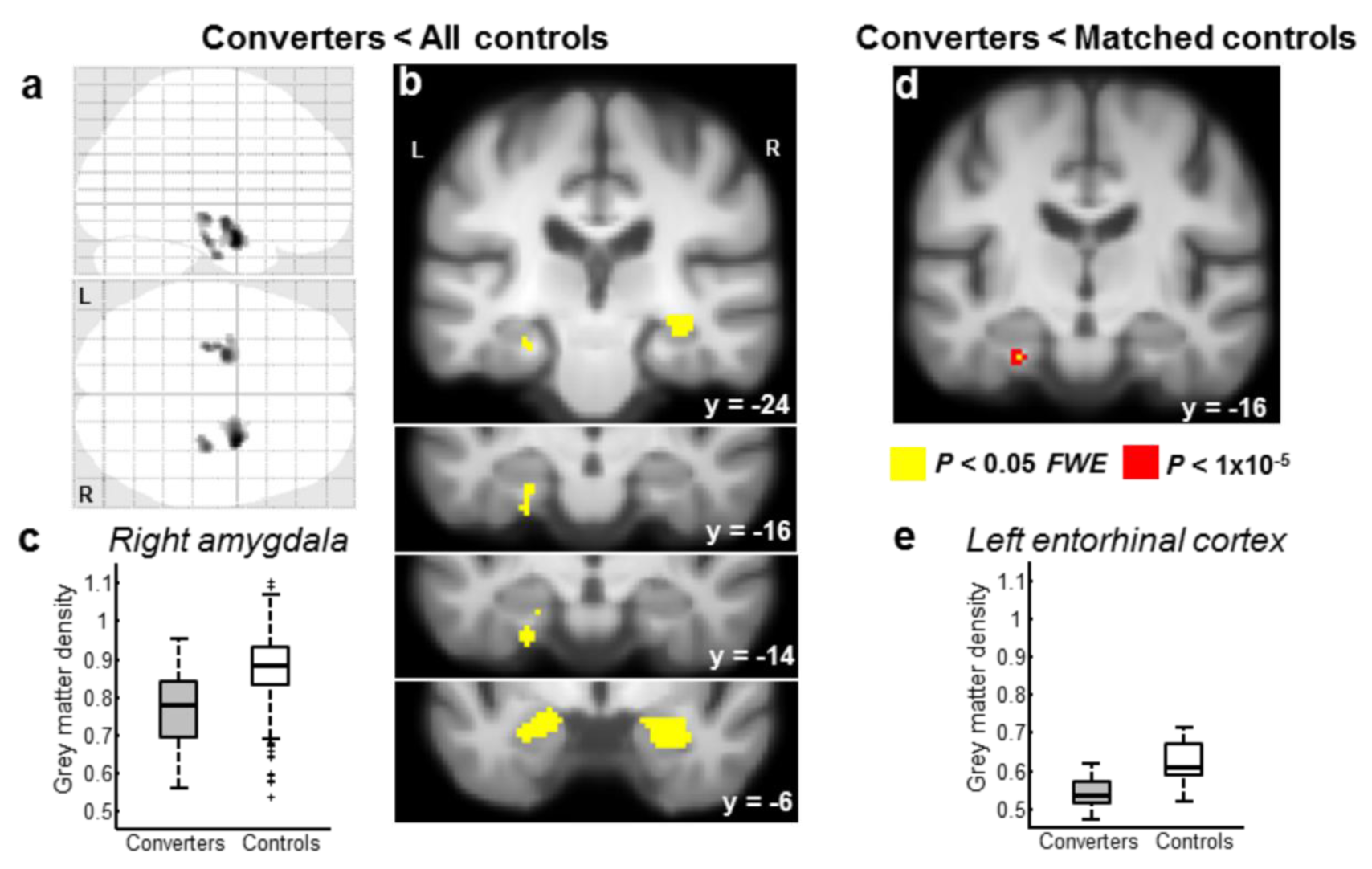
Medial temporal atrophy in healthy elderly individuals destined for MCI. Decreased grey matter density (GMD) in Visit 1 for 23 subsequent converters *vs*. all 790 controls (**a-c**) is limited to the medial temporal lobe. Reduced GMD, selective to bilateral amygdala, bilateral hippocampus, and left EC in converters one year before MCI onset, is depicted (**a**) on a glass-brain and (**b**) on serial coronal sections, overlaid on the group averaged T1 scan (threshold *P* < 0.05 whole-brain family-wise error corrected). (**c**) GMD in the global peak voxel (right amygdala; 30, - 3, −22) is plotted for converters and controls. For each boxplot (here and in Figure 3), the central mark indicates the median, and the bottom and top edges of the box indicate the 25th and 75th percentiles, respectively. Whiskers extend 1.5 times the interquartile range away from the top or bottom of the box, and outliers (+) are plotted individually. (**d**) In the comparison of the 23 converters *vs*. 23 matched controls, only left EC (−26, −16, −28) survives whole-brain family-wise error correction at *P* < 0.05. This effect is overlaid on the group average T1 scan in yellow (and at a more liberal threshold of *P* < 1×10^−5^ uncorrected in red) and GMD values plotted in (**e**).

The human EC has been segregated into posteromedial (pmEC) and anterolateral (alEC) portions on the basis of their patterns of functional connectivity^35-36^. To further refine the anatomical specificity of the EC GMD effect we observed, anatomical alEC and pmEC template images^36^ were warped to each participant’s anatomical image (**Supplementary Figure 1**). By averaging these warped templates over all participants, we obtained a template for each EC region for our study sample. The peak voxel within EC indexed by the comparison between converters and matched controls GMD (**Figure 2d-e**) localizes to alEC. However, to specifically test for differential reduction of GMD in anterolateral *vs*. posteromedial EC in converters *vs*. controls, we extracted the mean GMD from each region (alEC and pmEC), and hemisphere for each participant (**Supplementary Figure 1**). In the comparison of converters *vs*. matched controls, these were entered into a repeated measures ANOVA (with age, MMSE, years of education and TIV included as covariates). This analysis revealed a significant effect of group (*F*_1,_40 = 9.692, *P* = 0.003, η^2^_p_ = 0.195), a main effect of alEC *vs*. pmEC (*F*_1,40_ = 4.817, *P* = 0.034, η^2^_p_ = 0.107), and no main effect of hemisphere (*F*_1,40_ = 0.327, *P* = 0.570). The interaction between group and EC portion (*F*_1,40_ = 2.288, *P* = 0.138) and between group, hemisphere and EC portion (*F*_1,40_ = 1.523, *P* = 0.224) were not significant (**Supplementary Table 3**). Similar effects were obtained if converters were compared to all controls, and when these analyses were repeated on GMD images that had been spatially smoothed with a Gaussian kernel of 6mm at full-width half maximum (**Supplementary Table 3**). These results indicate that both portions of EC show reduced GMD one year prior to MCI diagnosis.

### The transition from healthy to MCI

We next examined the longitudinal trajectory of the converters relative to controls from V1 to V2. Specifically, we verified the likelihood that our cohort of converters was following decline compatible with AD neurodegeneration by examining differential atrophy rates from V1 to V2 in converters *vs*. controls. As shown in **Figure 3**, the pattern of atrophy in this one-year period that was significantly greater for the converter group *vs*. all non-converters is restricted to the medial temporal lobes bilaterally, in left amygdala extending into EC and in right EC extending into hippocampal body (**Supplementary Table 4**). Neuropsychological scores in this one-year time interval show significant worsening in MMSE, FCSRT and FAQ scores in converters relative to controls (**Supplementary Table 5**), consistent with development of MCI involving an amnestic component. White matter lesion load^34^ change during this 1-year interval showed no difference between converter and control groups (**Supplementary Table 5**).

**Figure 3.**
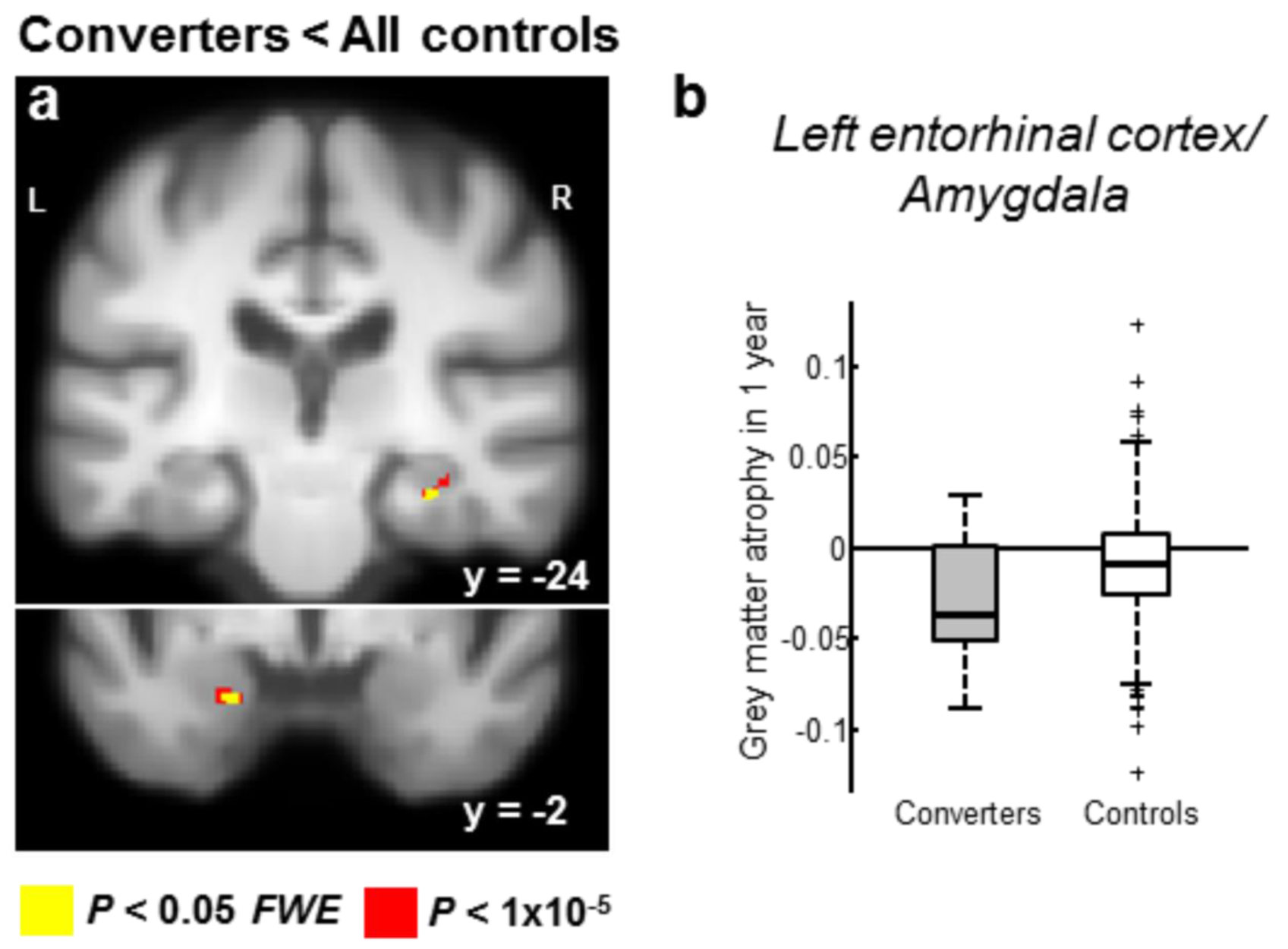
Atrophy rates in the medial temporal lobe over one year are significantly greater in those who develop MCI than those who remain cognitively intact. Accelerated grey matter atrophy, calculated from Visit 1 to 2, for 21 subsequent converters *vs*. 625 controls is limited to the medial temporal lobe. (**a**) Increased atrophy rates, selective to left amygdala/EC and right EC/hippocampus in converters is depicted on serial coronal sections, overlaid on the group averaged T1 scan (threshold *P* < 0.05 whole-brain family-wise error corrected in yellow and a more liberal threshold of *P* < 1×10^−5^ uncorrected in red). (**b**) Atrophy rates in the global peak voxel (left amygdala/EC; −21, −2, −26) are plotted for converters and controls.

### Predicting future MCI development

To explore the predictive performance of the different classes of data acquired in our cohort, we set up six different classification problems, calculating an elastic net regularized logistic regression on (i) demographic variables alone (age, gender, years of education), (ii) demographic variables plus APOEε4 genotype, (iii) neuropsychological variables alone (MMSE, FAQ, FCSRT, Rey–Osterrieth Complex Figure scores, Phonological Verbal Fluency, Semantic Verbal Fluency, State-Trait Anxiety Inventory (STAI)), (iv) demographic plus neuropsychological variables (that is, variables acquired without need for APOE genotyping or MRI scanning), (v) MRI-derived measures alone, which comprised hippocampal volumes and GMD values of 1248 2×2×2mm voxels from left and right entorhinal cortex (incorporated within the anterior parahippocampal gyrus mask of the FSL-Harvard-Oxford atlas), and (vi) all data modalities together. For this sixth classifier, demographic, neuropsychological and imaging variables (1262 variables in total) were included in the same model and entered into the elastic net estimator. For each classifier, we split the data at random into training/testing subsets using repeated 10-fold Cross-Validation (*i*.*e*., a data subset comprising 90% of the original data was used to train the regression model while 10% of the data was held-out to assess out-of-sample model prediction performance). Note that data from 42 control participants, chosen pseudorandomly, were not introduced into these models, so as to serve as “unseen” control participants in a subsequent independent validation of our model (described below; **Supplementary Table 6-7**). Thus 720 controls were used in initial classifier construction (**Figure 1)**.

The results of logistic regression with elastic net regularization on individual variable groups showed increasingly effective classification (as indexed by area-under-the-curve, AUC) in the order demographic < demographic plus APOEε4 genotype < MRI < neuropsychological parameters (**Figure 4a**). Adding demographic parameters to neuropsychological model did not improve performance, as the demographic variables were not selected as important for classification by the elastic net. However, when all modalities of data were used (All Modalities model), the ROC curve showed best performance, with best cross-validated AUC = 0.919 (performance measures for the 6 classifications, along with model coefficients, are provided in **Supplementary Table 8**). For the All Modalities model, out of 1262 possible variables, the elastic net selected only 3 coefficients. One was a neuropsychological variable, FCSRT delayed recall, which constituted the most important variable for classification. The remaining variables were GMD values for two voxels in left EC (**Supplementary Table 8**). Our MCI prediction equation when fit, (with all terms except the intercept ranked left to right in order of decreasing variable importance) is given in **equation (1)**.

**Figure 4.**
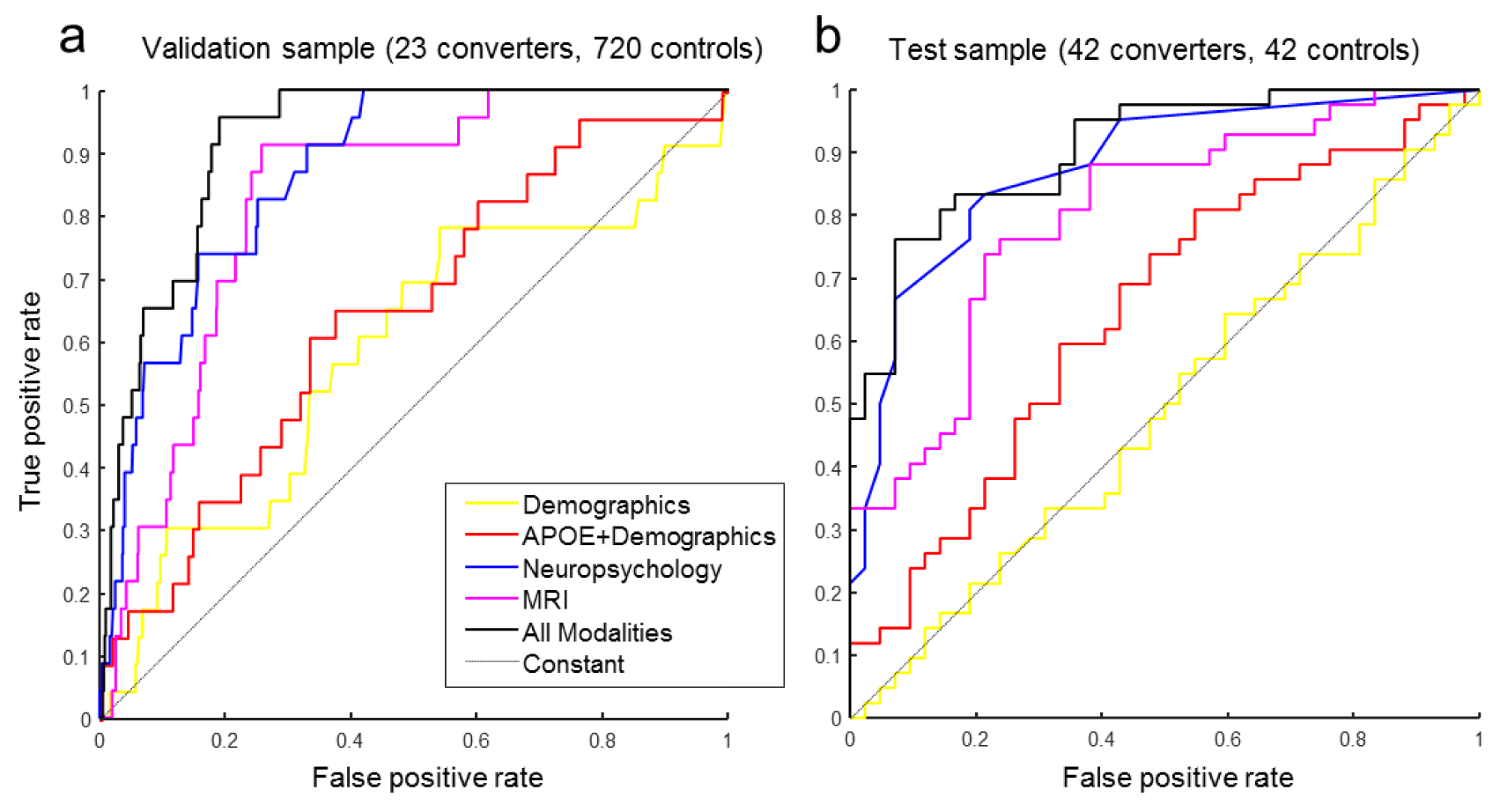
Predicting conversion to MCI within one-year. **a**. Receiver operating characteristic (ROC) curves generated by logistic regression with elastic net regularization on variables from visit 1 in our cohort of *n*=743 participants (23 subsequent converters and 720 controls). ROC curves are plotted for each group of variables individually: demographic variables (age, gender, years of education) (Demographics) and demographic variables plus APOEε4 genotype (APOE+Demographics), neuropsychological variables (Neuropsychology) alone (MMSE, FAQ, FCSRT, Rey–Osterrieth Complex figure, phonological verbal fluency, semantic verbal fluency, GDS and STAI), MRI-derived measures (MRI) alone (hippocampal volumes and GMD values of 1248 2×2×2mm voxels from left and right entorhinal cortex), and all data modalities combined (All Modalities). **b**. The ROC curve for the independent test cohort (unseen during training) is shown. Applying the All Modalities model to our new, independent sample of these new 42 converters and 42 controls yielded an Area-Under-the-Curve=0.905.

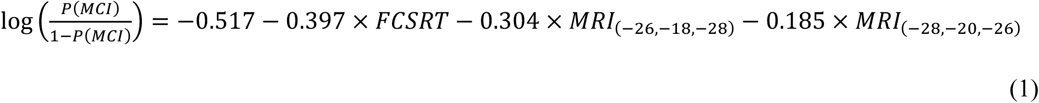

where *P(MCI)* is the probability of developing MCI within one year, *FCSRT* is total delayed recall performance on the FCSRT memory test, and *MRI*_(x, y, z)_ is the grey matter density at MNI coordinates *x, y, z* (both in left entorhinal cortex). The variables are assumed to be standardized, *i*.*e*., all variables are expected to be zero mean and unit variance, to render them (and their coefficients) comparable. The equations with coefficients for use with raw (not normalized) variables for this, the All Modalities model, as well as for the Neuropsychology model, are provided in **Supplementary equations (1) and (2)**.

Thus, an independent analytical approach – logistic regression with elastic net regularization – demonstrated that the same variables that ensued from our mass-univariate analyses performed best on distinguishing between subsequent converters *vs*. controls. Interestingly, age, gender and APOEε4 genotype, despite being known dementia risk factors, were not selected by the elastic net classifier as predictors for those destined for MCI at one-year.

### Test of the predictive model with unseen converters and controls

We next evaluated the classification performance of our trained statistical model on independent data derived from the Vallecas project. This project is currently in its 8^th^ year of yearly follow-up. We therefore interrogated all available data from all visits of the cohort to determine further individuals who transitioned from cognitively healthy to MCI over a 1 year time period. We denote the visit at which the MCI diagnosis is made as V_conv_, and the preceding visit (while still cognitively healthy) as V_conv-1_. Thus, on interrogation of the data from 2017-2020, we identified 42 “test” converters who converted on later visits of the Vallecas project. These individuals satisfied the same criteria used to define our original cohort of 23 V1 to V2 “original” converters, including the requirement for cognitive impairment to be present at the subsequent visit after conversion. Of the 42 converters, 17 developed amnestic MCI and 25 multi-domain MCI.

Baseline parameters at V_conv-1_ for these individuals who converted later in the Vallecas project are highly similar to those of the original V1 to V2 converter group (**Table 2**). As these converters were drawn from later visits of the Vallecas project, they are naturally older on average than our pool of V1 controls. Again, for test converters, APOEε4 and FCSRT showed robust effects relative to the entire (*i*.*e*., unmatched) control group, but for this comparison, FAQ, semantic verbal fluency and hippocampal volume also reached corrected significance. There were no significant differences between original V1 to V2 converters *vs*. test converters. Similarly, the change in cognitive function from V_conv-1_ to V_conv_ was significant in terms of MMSE, FAQ and FCSRT delayed total recall, with no difference observed between the original and test converters (**Supplementary Table 6**). In this older group of converters, white matter lesions (as measured by the Fazekas score^34^) also increased significantly in the year to conversion. The voxel-based morphometry comparison of test converters *vs*. controls showed a similar pattern of GMD reduction in the medial temporal lobes bilaterally (**Supplementary Table 7; Supplementary Figure 2**) with respect to the same comparison between original converters *vs*. unmatched controls (**Figure 2**), albeit more extensive in this older group of test converters.

The ROC curve for the different models derived from V1 to V2 converters in this independent cohort of 42 converters and 42 controls (that had been excluded during model training) is shown in **Figure 4b**. Classification performance is similar for validation and test samples for the 6 models. Critically, applying the classification model shown in **equation (1)** to this test sample yielded an AUC (±95% CI) = 0.905 (0.825 0.954), with specificity = 0.929 and sensitivity = 0.762. This remains the best performing model, although the AUC difference was not significantly different for the All Modalities *vs*. Neuropsychological model (**Supplementary Table 9**), the latter requiring only FCSRT and FAQ scores to reach test sample AUC (±95% CI) = 0.876 (0.783 0.938).

## DISCUSSION

We studied a large, single-site cohort, with yearly neuroimaging, and neuropsychological and clinical evaluation to determine biomarkers distinguishing between 2 groups of cognitively intact elderly subjects: those that develop MCI the following year and remain cognitively impaired in year 3 *vs*. those that remain cognitively healthy. Despite the two groups being both psychometrically within the normal ranges and clinically indistinguishable at year 1, we demonstrate highly selective differences – reduced delayed verbal memory scores, functional activities of daily life (FAQ), and left EC GMD – in healthy elderly subjects with impending MCI. This selectivity arises by comparing with a subgroup of controls chosen using exact matching for gender and APOE genotype (2 known risk factors for AD) and propensity score calculation for age, years of education, MMSE and TIV. By extracting a randomized experimental design from the full data-set, this procedure offers new and more robust interpretations of between-group differences, since known risk factors, such as APOEε4 genotype (more prevalent in the converter group), have also been shown to influence neuropsychological performance^26^ as well as brain structure^24^. In short, we can view with increased confidence reduced GMD in these localized areas as being causally related to the underlying mechanisms supporting the likelihood of developing MCI as opposed to merely demonstrating a weaker (probably confounded) association.

Neuropsychological evaluation revealed that although at baseline all converters had scores which fell entirely within the normal standard range of the control group on the different tests, delayed verbal memory (FCSRT) and FAQ were already significantly different between both groups. This suggests that delayed verbal memory recall (but not delayed visual memory assessed by the Rey–Osterrieth complex figure test) and functional activities can be considered neuropsychological markers in early, asymptomatic states of the disease. Indeed, in a large-scale population study^37^, FCSRT free recall in cognitively normal participants exhibited good sensitivity and fair specificity for AD prediction at 5 years (92% and 64% respectively), but showed poor positive predictive value (∼8%). The observation that FAQ scores differed between future converters and controls supports recent evidence that impairment in certain instrumental activities of daily living predicts greater risk of progressing from a diagnosis of cognitively normal to MCI^38^.

Differences in GMD between subsequent converters and controls were limited to MTL, with the pattern of atrophy in EC showing left-sided predominance, as has been reported in previous studies of MCI^39^. There is growing evidence that EC atrophy is a predictor for conversion to AD in patients with MCI^40-42^. Furthermore, in autosomal dominant, familial AD, longitudinal studies show significant atrophy in both EC and hippocampus 3.5 years before clinical diagnosis^21^. By contrast, large-scale observational studies of presymptomatic phases of sporadic AD are currently limited^13^. These studies^14-16^ involved smaller sample sizes than described here. Structural MRI data were acquired on 1.5T scanners^14-16,19^ as opposed to the 3T data we present, and they have used a region-of-interest approach, limiting comparisons to the medial temporal lobe^14-16^ with regions manually delineated^14,16^, as opposed to the whole-brain approach we report. Critically, previous studies have not used a matched-sampling framework to minimize bias introduced by background covariates that cannot be adequately controlled by simple linear adjustment and which can have profound implications on final inferential conclusions. Nevertheless, these studies also point towards EC as a brain region that is atrophied in asymptomatic elderly individuals destined for MCI, which is also supported by data from a small sample of healthy normal elderly showing that decreased glucose metabolism in EC is a predictor of impending MCI^43^. A larger prior study of progression of healthy to cognitive impairment^14^ (511 healthy individuals aged 60-90) performed manual region-of-interest (ROI) tracing on 1.5T scans and identified an association between reduction in hippocampal and amygdala volumes with the risk of developing of dementia, but did not measure EC volumes. Our data, however, show only a weak difference in hippocampal volume (automatic extraction at 3T) between future converters *vs*. non-converters. We note, however, that hippocampal atrophy indexed by our voxel-wise analysis (**Figure 2a, Supplementary Table 1**) is limited to anterior hippocampus, suggesting that volume measures subdivided along the hippocampal long-axis^44^ might reveal more pronounced differences in hippocampal head.

The ROI approach to EC volume calculation employed in prior studies^15-16^ does not consider recently observed functional subdivisions within this structure. Whereas the anatomical and functional dissociations of the lateral and medial entorhinal cortex in the rodent are well established, these two regions in the human have only recently been segregated into posteromedial and anterolateral portions on the basis of their patterns of functional connectivity^35-36^. In view of the suggested diverging cognitive roles of these two subregions of EC^35-36^, determining which area shows most atrophy prior to MCI development is important for predicting subtle cognitive deficits not immediately apparent in otherwise asymptomatic individuals destined for MCI. In rodents, the medial EC, the putative human homologue is the posteromedial part of EC^35^, is essential for spatial navigation^45^. However, we do not see differences in a direct test of GMD in the two EC portions, contrary to a previous suggestion that onset of MCI is in lateral EC^46^, when using functional measures of cerebral blood volume.

Having established (with a mass-univariate approach) biomarkers that distinguish between cognitively normal elderly individuals on the basis of progression to MCI within 1 year, we next explored the predictive capability of these parameters. This proceeded in an unbiased manner: all parameters were entered into a logistic regression with elastic net-regularization and automatic variable selection^29^. Consistent with our mass-univariate results, the combination of EC GMD and delayed verbal memory scores are most efficacious at predicting conversion from a cognitively healthy state to a state of MCI in this population. Our cross-validated model was effective in classifying a test sample of later converters and controls that were not used in the creation of the original model. Such out-of-sample testing is uncommon in previous studies on prediction of MCI in healthy elderly^18^. As a next step, it will be critical to test the generalizability of our statistical model when data from large-scale population studies employing 3T MRI scanning (*e*.*g*., the DELCODE study^47^) become available.

Both our original and test sample of converters comprised individuals who developed amnestic and multi-domain MCI. Despite this diagnostic heterogeneity, both subtypes show impaired episodic memory, which is seen most commonly in MCI patients who subsequently progress to a diagnosis of AD dementia^48^. That is, distinct MCI subtypes may have different outcomes, with the amnestic (single or multi-domain) MCI subtype being associated with a higher risk of progression to AD. This is important in the context of prediction, as in studies that mix amnestic and non-amestic future MCI converters, structural MRI parameters from prefrontal and parietal cortex show most discriminative power between converters and controls^49^, as opposed to the selective MTL effects we observe.

A potential limitation to our analyses is that the primary outcome of MCI was based only on clinical assessment, without identification of *in vivo* β-amyloid or tau, or post-mortem neuropathological diagnosis, which may have led to classification errors. MCI is a clinical and etiological heterogeneous entity, which can evolve to different dementia syndromes, remain stable or even revert to a state of normal cognition^50^. To mitigate classification errors, we integrated the clinical information from the visit after MCI diagnosis to exclude individuals who either reverted to a healthy state at or developed other neurological features outside of the most common MCI due to AD. The lack of a significant difference in neuroradiological index of white matter disease (Fazekas score) between V1 to V2 converters and controls indicates that a vascular cause for cognitive decline in the converters is less likely in this group of converters. Moreover, the observation of reduced GMD in EC of asymptomatic individuals destined for MCI is in keeping with the neuroanatomical pattern of post-mortem neurofibrillary changes characterizing Braak stage I-II, traditionally considered clinically silent^2,51^. Furthermore, whole-brain longitudinal analysis of atrophy from visit 1 to visit 2 revealed a focal decline in hippocampus and EC that accompanied conversion to MCI, indicating that most of the converters are likely marching towards an AD-type pathology.

FCSRT, FAQ and EC GMD measures can all be acquired non-invasively and relatively routinely in the clinical setting. The predictive algorithm presented here could therefore be implemented as a first screen to detect individuals destined to present with MCI the following year. Certain protective strategies could, at this stage, be implemented, such as blood pressure management^52-54^, management of depression and diabetes^7^, and exercise, smoking cessation and dietary advice^7^, potentially beneficial to elderly individuals at risk of MCI regardless of underlying pathology. This could be followed by measurement of brain, CSF or indeed plasma^55-56^ β-amyloid and tau levels, representing a step-wise strategy for screening to identify those at risk of impending decline in cognitive function specific to aetiology. Most importantly, at the time of identification, these individuals are functioning at a high level, allowing them to make decisions about their future care and treatment and make personal life choices at a time when it is still optimal to do so.

## Acknowledgements

We thank the participants of the Vallecas Project and the staff of the CIEN Foundation, and CJ Long for contributions to methodology employed here. This work was supported by the CIEN Foundation and the Queen Sofia Foundation, grants from Carlos III Institute of Health and Plan Nacional (SAF2016-78603-R) to MM, an Academy of Finland grant (316258) to JT and by a grant from the Alzheimer’s Association (2016-NIRG-397128) to BAS.

## Author contributions

B.A.S., J.T., L.Z. and A.S.-M. analysed the data. E.A. acquired imaging data and performed imaging quality control. M.M. supervised the project and organized data quality control. B.A.S. wrote the manuscript with input from all other authors.

## ONLINE METHODS

### Participants

Subjects involved in this study were volunteers participating in the ‘Alzheimer’s disease Vallecas Project’, a single-centre longitudinal community-based study^30^, currently in its eighth year. Participants, recruited by advertisement mostly from the Vallecas area in Madrid, Spain, provided written informed consent. The project was approved by the ethics committee of the Carlos III Institute of Health. From the initial pool of *n*=2077 contacted participants, the final sample size was *n*=1213 after excluding those that were not interested in participating in the study or met some of the exclusion criteria (**Figure 1**).

Inclusion criteria were as follows: 1) community-dwelling individuals; 2) both sexes; 3) from 69 to 86 years of age; 4) independent for activities of daily living; 5) no neurological or psychiatric disorder impeding daily functioning; 6) reasonable expectation of survival at a 4-year period, operationalized as absence of any severe disease at recruitment; and 7) able to sign informed consent. Exclusion criteria comprised: 1) dementia or severe cognitive deterioration, operationalized as Mini Mental Statement Examination (MMSE)^57^ below 24 and Functional Activities Questionnaire (FAQ)^58^ scores over 6 at the baseline assessment; 2) history of neurological or psychiatric disease with clinically relevant impact on cognition (*e*.*g*., cerebrovascular disease, major depression); 3) incidental structural brain findings with impact on cognitive impairment or survival (*e*.*g*., malignant brain tumour); 4) presence of a severe systemic disease (*e*.*g*., cancer under treatment); and 5) problems for understanding spoken or written Spanish language.

For the purpose of the current study, we applied further specific exclusion criteria. In our first analyses, conducted in 2015-2016, we identified individuals who converted from cognitively healthy to MCI from baseline visit (V1) to V2 **(Figure 1)**. To approximate cognitive normality at baseline, we selected participants with Clinical Dementia Rating (CDR)=0 or MMSE>26. We excluded 1) ‘reverters’ in V3 (namely participants who were considered converters in V2, but returned to a cognitive normal state in V3), 2) participants with incidental finding on MRI (such as large space-occupying lesions that invalidated the volumetric analysis), and 3) participants developing any other non-AD neurodegenerative disease in V2 or V3.

In addition to the 23 participants who converted to MCI on Visit 2 and fulfilled the above criteria, on subsequent interrogation of the data for the period 2017 to 2020, we identified a further 42 individuals who converted at later visits during the Vallecas project longitudinal study (4 more from V1 to V2, 10 from V2 to V3, 11 from V3 to V4, 4 from V4 to V5, 5 from V5 to V6, and 8 from V6 to V7). As for the original cohort of converters, all participants had Clinical Dementia Rating (CDR)=0 and MMSE>26 on the visit prior to conversion. We again excluded those participants whom, on the visit after being diagnosed with MCI, developed any other non-AD neurodegenerative disease or reverted to a cognitively healthy state.

### Yearly evaluation

In the baseline visit, sociodemographic data, vital signs, and blood samples (for measuring *APOE* genotype) were collected, followed by neuropsychological, clinical, and multi-sequence MRI assessment. In the following visits in subsequent years (V2 onwards) the same procedure was repeated, excluding genetic testing. Neuropsychological testing comprised a comprehensive battery including the following tests: *Cognitive performance:* MMSE, FCSRT, Rey–Osterrieth Complex Figure (acquired in all visits except V2) and phonological and semantic verbal fluency; *Depression and Anxiety:* Geriatric Depression Scale, State-Trait Anxiety Inventory; *Functional scales:* CDR, FAQ. A more detailed description of the Vallecas Project design, demographic and neuropsychological measures and clinical assessments is described elsewhere^30,59^.

A diagnosis of MCI was made when the following criteria were fulfilled^60^ (1) concern regarding a change in cognition, from the patient, a proxy informant or a trained clinician, (2) impairment in one or more cognitive domain (performance is typically below 1-1.5 SD, according to participant age and education, but these ranges are guidelines and not cut-off scores), (3) preservation of independence in functional abilities, (4) not demented. MCI diagnosis was further split into the three subgroups of amnestic, non-amnestic and mixed^61^. Participants who developed MCI in the follow-up visits were considered ‘converters’ and those who remained cognitively healthy were considered ‘controls’. The diagnosis of MCI was agreed between 2 experienced clinicians, one neurologist and one neuropsychologist. In the case of lack of agreement between neurologist and neuropsychologist about the diagnosis of particular individual, the case was reviewed at an independent consensus meeting involving 3 further members of the research team (neurologists and neuropsychologists). Importantly, at the time of making the diagnosis of MCI, the clinical team was blind to neuropsychological test scores obtained the previous year (*i*.*e*., those used in the predictive algorithm), as well as to the current MRI scanning. That is, all the diagnoses were made only with the clinical information available at each visit and without knowing any detail about the cognitive trajectory of the participants, precluding a circularity bias.

### APOE genotyping

Total DNA was isolated from peripheral blood following standard procedures. Genotyping of APOE polymorphisms (rs429358 and rs7412) was determined by Real-Time PCR^62^. Failure rate of genotyping was 0.3%. The frequency of APOE ε4 allele in our cohort is 17.6%, consistent with previous findings in the Spanish population^63^.

### Matched sampling

Because our grouping (MCI conversion) variable lies outside of experimental control, inference will be biased due to two interrelated factors (a) parametric model misspecification, and (b) treatment-control group covariate imbalance. While in experimental design, randomization provides some guarantee that treatment and control groups are only randomly different in background attributes, valid inference in the present scenario requires the extraction of a randomized design (assuming one exists) from the dataset. To this end, and to mitigate the effect of potential confounds, we used the framework of potential outcomes to develop a matched-sampling procedure^64^. The potential outcomes approach postulates a counterfactual model for subject *i* in the conversion group and seeks to estimate how the measured outcome for subject *i* would have been manifested had they not undergone the conversion. Matching, in its simplest form, estimates the unobserved counterfactual by selecting an observed outcome measure from the potential control group that is an exact match in all of the measured background variables. However, in practice, there is a curse of dimensionality problem when the number of discrete background covariates is either large relative to the number of subjects, or when one or more of the balancing covariates contains continuous values. In either case, approximate methods are necessary which can summarize, while preserving certain key properties, large numbers of covariates into a convenient one dimensional summary. The propensity score is one such measure and is defined as the probability of a subject being classified as an MCI converter conditional on everything that is known *a priori* about that subject that does not influence either the likelihood of MCI or the outcome variable. The propensity score is known as an equal percent bias reducing (EPBR) technique^64^ meaning that if a close match is obtained in the propensity score distribution between the groups then the groups will also be close in the original covariates. For this study we implemented a two-level optimization procedure that first ranks covariates by importance followed by exact matching on the main risk factor variables associated with AD: gender and APOEε4 status. The exact matching yields subgroups of subjects identical in these risk factors. Within each of the 23 subgroups (one subgroup per member in the converter group), the subjects were next assigned a propensity-based distance relative to their assigned converter. Next within each subgroup each potential control is ranked according to the propensity score distance computed from age, years of education, MMSE and total intracranial head volume at Visit 1. We next need to decide *k*, the top number of controls to be selected as the optimal control group. Each of our 23 converter subgroups had at least one potential control and so we chose *k*=1, corresponding to a pair-matched experimental design. While this choice implies we have minimized bias with some correspondent increase in variance, we have improved the interpretability of our subsequent outcome measures. Our effect-sizes in this particular application are strong relative to variance reduction of bias, which, in our opinion, dominates this choice over a variable-*k* matching. For the matching procedure, we performed list-wise deletion of participants whose data in one or more demographic or neuropsychological variables at V1 were missing, leading to a further 26 control participants being excluded. We note that list-wise deletion is known to incur a potentially significant estimation bias depending on the nature and scale of subject attrition^65^.

### Statistical analyses: Neuropsychological, genetic and demographic group comparisons

These variables were analysed using SPSS software (version 21.0; SPSS Inc., Chicago, USA). Welch’s t-test and χ2 test were used to compare quantitative and qualitative variables respectively, between converters and controls in V1 and subsequently for test converters in V_conv-1_. A two-way analysis of variance (ANOVA) was performed for testing the interaction of the different neuropsychological measures in converters and controls between V_conv-1_ and V_conv_.

### Brain Imaging

#### Image acquisition

All magnetic resonance images were acquired using the 3 Tesla MRI (Signa HDxt General Electric, Waukesha, USA) at the Queen Sofia Foundation for Alzheimer’s Research, Madrid Spain. Using a proprietary phased array 8 channel head coil, whole brain T1-weighted images were acquired for each participant using the following protocol: 3D sagittal sequence fast spoiled gradient recalled (FSPGR) with inversion recovery (repetition/echo/inversion time 10/4.5/600ms, field-of-view=240mm, matrix=288×288, 166 sagittal slices of thickness=1mm), yielding an overall non-isotropic image resolution of 1.0×0.5×0.5mm.

#### Fazekas scoring

This was performed by a neuroradiologist and recorded as the higher value of periventricular or deep white matter hyperintensities score, assessed using fluid-attenuated inversion recovery (FLAIR) imaging (repetition/echo/inversion time 9000/130/2100ms, field-of-view=240mm, slice thickness 3.4cm).

#### Cross-sectional grey matter density analysis on Visit 1

Voxel-based morphometry (VBM) analysis^66^, using the DARTEL (Diffeomorphic Atlas Registration Tool with Exponentiated Lie Algebras) suite within statistical parametric mapping SPM12 software (Welcome Trust Centre for Neuroimaging, University College London; http://www.fil.ion.ucl.ac.uk/spm/), was performed to compare whole-brain GMD between converters and controls on V1. For each participant, the T1-weighted structural image was first registered to a common MNI anatomical orientation, using a low dimensional affine transformation, bias corrected to mitigate potential inhomogeneities in the image intensities and resliced to 1mm isotropic resolution. Images were then segmented into grey matter, white matter and CSF. A nonlinear spatial registration technique (DARTEL) was then applied to the grey matter tissue maps. The template for registration was constructed using all participants from our converter group plus selected control participants. For each V1 to V2 converter, those participants from the control group who were exact matches in terms of age (in years), gender and total intracranial volume (discretized after computing the bin width based on the data interquartile range bin-size=2*(Q_3_-Q_1_)*n^-1/3^, into four categories), were selected. This resulted in a total of 348 participants (all converters plus 325 controls) being included in the DARTEL template creation.

Next, each participant’s segmented grey matter map was “modulated” by the Jacobian map to preserve the amount of grey matter signal relative to the original (unwarped) image. Finally, each modulated gray matter map was affine transformed onto the MNI template and smoothed with a Gaussian kernel of 6-mm full width at half maximum. These images were next entered into a two-sample *t*-test, comparing whole-brain differences in GMD of converters and controls. We first performed an analysis comparing converters *vs*. all controls, including as covariates of no interest age, gender, APOEε4, MMSE, years of education, and individual TIV values. The latter were obtained by summing the volumes of the grey matter, white matter and cerebrospinal fluid. Second, we repeated the same analysis entering converters and only matched controls. Age, MMSE, years of education and TIV values were again included as covariates of no interest, to account for residual imbalance following propensity score matching. There is currently a lack of consensus on whether to perform a paired *t*-test over two-sample *t*-test on matched data^67^. When doing a two sample *t*-test, it is assumed that the two samples are not dependent on each other. This assumption will not be violated just because the participants had similar demographics. That is, we do not view these as paired observations; we have only made the two populations more comparable by matching.

#### Entorhinal Cortex Anatomy

alEC and pmEC and masks are predicted clusters derived by multivariate classification of perirhinal cortex and parahippocampal cortex connectivity preference^36^. The construction of participant-specific masks for alEC and pmEC proceeded as follows. The high-resolution whole brain T1-weighted template (0.6 mm isotropic resolution) associated with the EC masks^36^ was segmented in SPM12 and the resultant grey matter image was normalized to the unsmoothed, modulated grey matter density image for each participant separately. The warp parameters ensuing from normalization were applied to the EC masks in order to obtain participant-specific masks of alEC and pmEC. The mean GMD from each EC subregion was then extracted.

#### Longitudinal analysis

Longitudinal registration, tissue segmentation, and spatial normalization were performed using the SPM12 pairwise longitudinal toolbox, which uses the time between scans to perform a “symmetric” registration of longitudinal scans to an estimated midpoint image. In addition to alleviating potential bias in the choice of reference image, this procedure ensures intra-participant images have identical processing to render their comparability across time. From the 23 converters, 21 had MRI studies in V2 as well as V1. Thus, T1 images from V1 and V2 were co-registered and the midpoint image calculated, as well as a map of the rate of volumetric change (the divergence field) estimated using the difference in warp fields of the two scans relative to the midpoint in unit time. The former serve as inputs for tissue segmentation, while the divergence field represent annualized volume change within participant. Thus, for each participant, the midpoint average image was segmented into grey matter, white matter and CSF and the ensuing grey matter image was multiplied by the divergence field to yield images of yearly grey matter atrophy. A two-sample *t*-test was performed on spatially normalized atrophy maps smoothed with a 6mm kernel comparing whole-brain differences between converters and controls.

#### Automated hippocampal volume extraction

Automatic segmentation of hippocampal subfields was performed on each participant’s T1-weighted image using FreeSurfer 5.3.0 (https://surfer.nmr.mgh.harvard.edu/). Eight subregions were obtained: CA1, CA2–3, CA4-Dentate gyrus, pre-subiculum, subiculum, fimbria, hippocampal tail, and hippocampal fissure. Segmentations for all participants were visually inspected. The whole hippocampus volume was obtained by adding subfields CA1, CA2–3, CA4-Dentate gyrus, hippocampal tail, and subiculum.

### Elastic-Net-penalized Logistic Regression for the Prediction of MCI

We set up six different classification problems and for each fit a logistic regression classifier with an elastic net penalty, implemented using the Matlab R2018b and glmnet toolbox^68^ (version 11 March 2015). The EC voxels entered into two of these classification problems were extracted from each participants’ grey matter density image (modulated, normalized to MNI space and smoothed at 6mm in SPM12 version v6225), with anatomical boundaries defined by the anterior parahippocampal gyrus mask of the FSL-Harvard-Oxford atlas (http://www.fmrib.ox.ac.uk/fsl/). This mask shows good overlap with EC in our elderly T1 scans, with the anterior extent including putative transentorhinal cortex. Before fitting the elastic net model, the data were standardized so that each variable had zero mean and unit variance. This was done without any reference to class labels. The missing data was imputed using k-nearest neighbours variable imputation^69^, with k = 3, again using no label information and imputing test data (42 test converters and matched controls) only based on the data from training and validation set (23 V2 converters and 720 controls). Note that in view of class imbalance, each member of the class with more samples was downweighted according to its empirical frequency. That is, we probability weighted the subjects depending on the frequency of each group (in a way that the weights in both groups sum to 0.5).

The elastic net penalty compensates for the large number of variables relative to number of subjects. This is achieved by combining a lasso (L1) penalty that performs an automatic model selection, often choosing groups of parameters that are most correlated, with regularization of the large number of potential variables by shrinking similar variables towards one another (the grouping property)^29^. Two hyperparameters need to be chosen in elastic net regression problems. Lambda controls the extent to which the model is penalized; a Lambda of 0 reduces the estimate to an unpenalised and perhaps unidentifiable ordinary least squares estimate whereas a large Lambda will result in a heavily penalized model with small coefficient vectors. The second hyperparameter is alpha, which governs the amount of interplay within the penalty, between the model selection and the assumed correlation between the parameters. A choice of alpha close to 1 will tend to focus on a highly sparse model selection and will tend to ignore potential groupings between the parameters. Conversely, a choice of alpha close to zero will lead to the inclusion of all parameters and will tend to exploit correlations across a larger number of parameters.

#### Model Fitting and Hyper-parameter Estimation

To select hyper-parameters that optimize model performance while protecting against overfitting, we performed a two-dimensional tuning grid search embedded within a cross-validation procedure. We chose lambda to run between 0 and 0.5 and alpha to run between 0 and 1 with a respective spacing of 0.01 resolution across the grid.

Model performance may be assessed using one of a number of metrics, such as R^2^, accuracy or Area under the curve (AUC). In our case we chose AUC as it was the most appropriate given our final study objectives. AUC can be taken as an estimate of the probability of the classifier ranking a randomly chosen positive example (converter) higher than a randomly chosen negative example (non-converter control). For each possible choice of alpha and lambda, we assessed model performance in the following manner.

First we split the data at random into training/testing subsets using repeated *k*-fold Cross-Validation with *k*=10. The cross-validation was stratified, *i*.*e*., each fold contained (approximately) equal number of converters. In standard *k*-fold cross-validation, uncertainty in performance estimates may be reduced by repeating each *k*-fold cross-validation *L* times (in our case *L=25*) and averaging across the *L* estimates returned from each single *k*-fold. Within each of the *k*-folds, one repeat iteration proceeded as follows: we fit the model to the training subset, and estimated the out-of-sample AUC from the held-out 10% subset. Next the held out data was returned to the main dataset and the process repeated until each subset of the data had been used in both model training and in assessing its out-of-sample performance on the unseen data subset. The AUC for a single cross-validation run was computed using the pooling method^70^ and the process iterated another *L-1* times. After the *L*^th^ repeat, the AUC measure was averaged over the *L* repeats onto the location on the tuning grid corresponding to that choice of alpha and lambda. The net result was a set of cross-validation error curves and their associated standard errors plotted as a function of tuning parameter. We next selected the point on these curves yielding maximal model performance (AUC) and plotted the corresponding (cross-validated) receiver operating characteristic (ROC) for elastic-net classifier with the optimal hyper-parameters for each of the data classification problems shown in **Figure 4a**. Confidence intervals for AUCs were computed with accelerated bias corrected bootstrap method implemented in the Matlab function *perfcurve* (Version 2018b). The DeLong test^71^ was used to derive *P*-values for the differences between different models (employed in the StaR online tool^72^).

#### Out-of-sample testing

To select a control group for the test converters (described in the section ‘**Participants**’ above), for each converter on visit V_conv-1_ (*i*.*e*., the data is from visit V_conv-1_, conversion at V_conv_), a control was randomly selected from visit V_conv-1_, with the constraints of being of the same age and gender as that converter. This control was then removed from the large set of controls. This process was continued until all converters had a matched control. The model with the optimal hyper-parameters, now trained with the combined training and validation set, was then subjected to testing with this out-of-sample test set of 42 converters and 42 matched controls. The ROC curve for this classification problem is plotted in **Figure 4b**. The *P*-values and CIs for AUCs were computed as described above for validation.

In calculating specificity and sensitivity for the test sample, we assumed that there is a dataset shift (due to learning effects on repeat neuropsychological tests over visits), hence, we cannot expect 0.5 threshold to be correct. We therefore derived a new optimal threshold/constant term by two methods. In an optimal threshold approach, used for reporting accuracy, sensitivity and specificity in the main manuscript for the All Modalities model, the labels in the test set were used to decide which cut-off threshold (converters *vs*. non-converters) is optimal in terms of (balanced) accuracy along the ROC curve (*i*.*e*., which point of ROC curve gives the best accuracy). Since the labels in the test set are being used, these thresholds are decided based on the test data. In the split-half method, motivated by ^73^, we sampled half of the test subjects (21 controls, 21 converters) and solved the optimal threshold (cut-off point) using (0,1)-criterion^74^. We then applied this threshold to calculate sensitivity and specificity values for the other half of subjects (the other 21 controls and 21 converters). Then, we reversed the roles of two split-halves and averaged resulting sensitivity and specificity values. This method produces two thresholds for the test sample, but it is a straight-forward way to calibrate the data for learning effects that can work with a small calibration set. Note that just using the optimal thresholds would lead to training on the test data type problem and tuning the decision threshold with linear classifiers is equivalent to tuning the constant parameter of the classifier. In reporting the results, we repeated split-half division 50 times to ensure that the results did not depend on a particular split-half division. Performance measures pertaining to both these optimal threshold approaches, and taking a threshold of 0.5, are provided in **Supplementary Table 8**.

## Data availability

The data that support the findings of this study are available from the corresponding author upon reasonable request.

## Code availability

Structural MRI data were analysed using SPM12 (http://www.fil.ion.ucl.ac.uk/spm) run on Matlab (The Mathworks). Cross-sectional and longitudinal analyses of V1 to V2 converters, conducted in 2015-2016, were performed using SPM12 beta version v6015. All subsequent analyses were performed with SPM12 version v6225. Propensity scoring was calculated within the R-environment for statistical computing (https://www.r-project.org/; using CRAN packages optmatch). Elastic net-regularized logistic regression was performed using the glmnet toolbox in Matlab. The KNN imputation Matlab-code is available at https://github.com/jussitohka/MCIpredict as well as the Matlab scripts to run the elastic net training and classification.

