## Supplementary Material for "Predicting the future development of mild cognitive impairment in the cognitively healthy elderly"

**Supplementary Table 1.** Local maxima for the group level comparison of grey matter density in 790 controls *vs*. 23 converters at visit 1 surviving whole-brain family-wise error (FWE) correction at $P<0.05$ (independent t-test with age, gender, years of education, MMSE, APOEε4 genotype and TIV included as covariates).

| Brain Area | x | y | z | *Z*-score | *P*-value (FWE-corr) |
|---|---|---|---|---|---|
| Right amygdala | 30 | -3 | -22 | 6.00 | 0.0001 |
| Left amygdala and anterior hippocampus | -20 | -10 | -12 | 5.68 | 0.0009 |
| Right hippocampal body | 34 | -21 | -10 | 5.54 | 0.0019 |
| Left posteromedial entorhinal cortex | -26 | -20 | -24 | 5.50 | 0.0023 |
| Left entorhinal cortex, extending laterally into transentorhinal cortex | -26 | -14 | -32 | 5.43 | 0.0032 |

**Supplementary Table 2.** Local maxima for the group level comparison of grey matter density in 23 matched controls *vs*. 23 converters at visit 1 surviving whole-brain family-wise error (FWE) correction at $P<0.05$ (independent t-test with age, MMSE, years of education and TIV included as covariate).

| Brain Area | x | y | z | *Z*-score | *P*-value (FWE-corr) |
|---|---|---|---|---|---|
| Left anterolateral entorhinal cortex | -26 | -16 | -28 | 5.14 | 0.035 |

**Supplementary Table 3.** Results of the repeated measures ANOVA of mean grey matter density in entorhinal cortex anterolateral (al) and posteromedial (pm) portions 23 converters to 790 controls at visit 1 (Unmatched) and 23 converters to 23 matched controls (Matched). For Unmatched comparisons, age, gender, years of education, MMSE, APOEε4 genotype and TIV were included as covariates. For Matched comparisons, age, MMSE, years of education and TIV were included as covariates in repeated measures ANOVAs and one-way ANOVAs. Significant *P* values and associated effect sizes are in bold.

| Comparison | *F* value | df | *P* value | *Partial eta-squared* |
|---|---|---|---|---|
| **Converters *vs.* Unmatched controls (no spatial smoothing)** | | | | |
| Effect of group | 20.537 | 1,805 | **<0.001** | **0.025** |
| Main effect of EC portion (al *vs.* pm) | 2.934 | 1,805 | 0.087 | 0.004 |
| Main effect of hemisphere | 0.006 | 1,805 | 0.938 | 0.000 |
| Interaction group by EC portion | 0.706 | 1,805 | 0.401 | 0.001 |
| Interaction group by EC portion by hemisphere | 2.186 | 1,805 | 0.140 | 0.003 |
| **Converters *vs.* Matched controls (no spatial smoothing)** | | | | |
| Effect of group | 11.594 | 1,40 | **0.002** | **0.225** |
| Main effect of EC portion (al *vs.* pm) | 4.217 | 1,40 | **0.047** | **0.095** |
| Main effect of hemisphere | 0.380 | 1,40 | 0.541 | 0.009 |
| Interaction group by EC portion | 3.352 | 1,40 | 0.075 | 0.077 |
| Interaction group by EC portion by hemisphere | 3.914 | 1,40 | 0.055 | 0.089 |
| **Converters *vs.* Unmatched controls (smoothing 6 mm)** | | | | |
| Effect of group | 25.002 | 1,805 | **<0.001** | **0.030** |
| Main effect of EC portion (al *vs.* pm) | 0.530 | 1,805 | 0.467 | 0.001 |
| Main effect of hemisphere | 0.225 | 1,805 | 0.635 | 0.000 |
| Interaction group by EC portion | 0.029 | 1,805 | 0.864 | 0.000 |
| Interaction group by EC portion by hemisphere | 0.244 | 1,805 | 0.622 | 0.000 |
| **Converters *vs.* Matched controls (smoothing 6 mm)** | | | | |
| Effect of group | 14.889 | 1,40 | **<0.001** | **0.271** |
| Main effect of EC portion (al *vs.* pm) | 3.197 | 1,40 | 0.081 | 0.074 |
| Main effect of hemisphere | 1.064 | 1,40 | 0.308 | 0.026 |
| Interaction group by EC portion | 2.151 | 1,40 | 0.150 | 0.051 |
| Interaction group by EC portion by hemisphere | 0.442 | 1,40 | 0.510 | 0.011 |

**Supplementary Table 4.** Local maxima for the group level comparison of grey matter atrophy (longitudinal VBM) in 625 controls *vs*. 21 converters from visit 1 to 2 surviving family-wise error (FWE) correction at *P*<0.05 (independent t-test with age, gender, years of education, MMSE, APOEε4 genotype and TIV at visit 1 included as covariates).

| Brain Area | x | y | z | *Z*-score | *P*-value (FWE-corr) |
|---|---|---|---|---|---|
| Left amygdala extending into entorhinal cortex | -21 | -2 | -26 | 4.88 | 0.0153 |
| Right entorhinal cortex extending into hippocampal body | 32 | -24 | -20 | 4.81 | 0.0208 |

**Supplementary Table 5.** Participant changes in neuropsychological scores from Visit 1 (V1) to Visit 2 (V2). Significant p-values are given in bold. *Survives Bonferroni correction for the 8 tests.

| | **Mean Difference V2-V1 in 21 Converters** | **Mean Difference V2-V1 All 625 Controls** | **Interaction (Difference V2 vs V1) x (Converter vs All Controls)**<br><br>ANOVA (F value and p value) |
|---|---|---|---|
| MMSE | 27.1 – 28.6 = -1.5 | 28.9 – 29.0 = -0.10 | *F* = 17.81<br>***P* = 2.8 x $10^{-5}$ *** |
| FAQ | 2.1 – 1.2 = 0.9 | 0.3 – 0.3 = 0.0 | *F* = 24.73<br>***P* = 8.5 x $10^{-7}$ *** |
| FCSRT delayed total recall | 10.4 – 11.8= -1.4 | 14.90 – 14.70 = 0.20 | *F* = 18.14<br>***P* = 2.4 x $10^{-5}$ *** |
| Phonological Verbal Fluency | 12.2 – 12.8 = -0.6 | 15.0 – 14.1 = 0.9 | *F* = 2.69<br>*P* = 0.10 |
| Semantic Verbal Fluency | 13.9 – 16.6 = -2.7 | 18.6–19.2 = -0.60 | *F* = 3.85<br>*P* = 0.0501 |
| GDS | 2.7 – 2.3 = 0.4 | 1.4 – 1.4 = 0.00 | *F* = 0.70<br>*P* = 0.40 |
| STAI | 17.4 – 17.2 = 0.2 | 16.70 – 16.80 = -0.10 | *F* = 0.09<br>*P* = 0.76 |
| Fazekas score | 1.14 – 1.10 = 0.04 | 1.14 – 1.11 = 0.03 | *F* = 0.06<br>*P* = 0.81 |
| CDR | 21: 0.5 (all of them diagnosed with MCI in V2) | 625: 0 | |

**Supplementary Table 6.** Test converter changes in neuropsychological scores from preconversion visit to conversion visit ($V_{conv-1}$ and $V_{conv}$). Note that the reduction in size of the control group (now 600) reflects the fact that some validation converters were originally controls at V1 and V2 (their data have been excluded to preclude mixing between- and within-subject comparisons). Significant *P*-values are given in bold. *Survives Bonferroni correction for the 8 tests.

| | **Mean Difference Post- vs Pre-conversion in 42 Test Converters** | **Mean Difference V2-V1 All 600 Controls** | **Interaction (Difference Post- *vs* Pre-conversion) x (Test Converter *vs* Controls)**<br><br>ANOVA (F value and p value) | **Interaction (Difference Post- *vs* Pre-conversion) x (Test Converter *vs* V1 to V2 Converter)**<br>ANOVA (F value and p value) |
|---|---|---|---|---|
| MMSE | 27.0 – 28.4 = -1.4 | 28.9 – 29.1 = -0.2 | *F* = 28.63<br>***P* = 1.2 x $10^{-7}$ *** | *F* = 0.02<br>*P* = 0.88 |
| FAQ | 2.5 – 1.2 = 1.3 | 0.33 – 0.32 = 0.01 | *F* = 90.67<br>***P* = 3.7 x $10^{-20}$ *** | *F* = 0.82<br>*P* = 0.37 |
| FCSRT delayed total recall | 11.2 – 13.4 = -2.2 | 15.0 – 14.7 = 0.3 | *F* = 110.93<br>***P* = 5.0 x $10^{-24}$ *** | *F* = 0.18<br>*P* = 0.14 |
| Phonological Verbal Fluency | 13.2 – 13.9 = -0.7 | 14.9 – 14.1 = 0.8 | *F* = 5.96<br>***P* = 0.015** | *F* = 0.05<br>*P* = 0.82 |
| Semantic Verbal Fluency | 13.9 – 15.9 = -2.0 | 18.5 –19.3 = -0.8 | *F* = 3.26<br>*P* = 0.072 | *F* = 0.62<br>*P* = 0.43 |
| GDS | 2.8 – 2.4 = 0.4 | 1.4 – 1.4 = 0.00 | *F* = 1.48<br>*P* = 0.22 | *F* = 0.01<br>*P* = 0.94 |
| STAI | 15.9 – 14.8 = 1.1 | 16.6 – 16.8 = -0.2 | *F* = 1.51<br>*P* = 0.22 | *F* = 0.11<br>*P* = 0.74 |
| Fazekas score | 1.47 – 1.33 = 0.14 | 1.13 – 1.10 = 0.03 | *F* = 8.42<br>***P* = 0.0038 *** | *F* = 6.57<br>***P* = 0.013** |
| CDR | 42: 0.5 | 600: 0 | | |

**Supplementary Table 7.** Local maxima for the group level comparison of grey matter density in 762 controls at visit 1 *vs.* 42 test converters at preconversion visit ($V_{conv-1}$) surviving whole-brain family-wise error (FWE) correction at $P$<0.05 (independent t-test with age, gender, years of education, MMSE, APOEε4 genotype and TIV included as covariates).

| **Brain Area** | **x** | **y** | **z** | ***Z*-score** | ***P*-value (FWE-corr)** |
|---|---|---|---|---|---|
| Left entorhinal cortex extending into parahippocampal gyrus | -26 | -24 | -22 | 6.98 | $5.49 \times 10^{-7}$ |
| Left anterior hippocampus extending into amygdala | -26 | -10 | -16 | 6.59 | $6.91 \times 10^{-6}$ |
| Left perirhinal cortex | -36 | -12 | -32 | 6.32 | $3.65 \times 10^{-5}$ |
| Right posterior hippocampus | 34 | -33 | -2 | 6.26 | $5.19 \times 10^{-5}$ |
| Right anterior hippocampus | 28 | -15 | -16 | 5.94 | $3.31 \times 10^{-4}$ |
| Right posterior hippocampus | 26 | -33 | -4 | 5.38 | 0.0066 |
| Right inferior temporal gyrus | 46 | -41 | -22 | 6.21 | $7.15 \times 10^{-5}$ |
| Left anterior cingulate gyrus | -8 | 49 | 7 | 6.02 | $2.10 \times 10^{-4}$ |
| Left ventromedial prefrontal cortex | -6 | 41 | -8 | 5.63 | 0.0018 |
| Left anterior cingulate gyrus | -10 | 47 | 0 | 5.49 | 0.0036 |
| Right superior temporal sulcus | 50 | 5 | -22 | 5.60 | 0.0021 |
| Right superior temporal sulcus | 56 | -5 | -16 | 5.50 | 0.0035 |
| Right temporal pole | 44 | 10 | -30 | 5.45 | 0.0046 |

**Supplementary Table 8.** 1-year conversion prediction results. Classification performance measures for each group of variables individually: demographic variables (age, gender, years of education) (Demographics) and demographic variables plus APOEε4 genotype (Demographics plus APOE genotype), neuropsychological variables (Neuropsychology) alone (MMSE, FAQ, FCSRT, Rey–Osterrieth Complex figure scores, phonological verbal fluency, semantic verbal fluency, and STAI), Neuropsychology plus Demographic variables, MRI-derived measures (MRI) alone (hippocampal volumes and GMD values of 1248 2x2x2mm voxels from left and right entorhinal cortex), and all data modalities combined (All Modalities). GMD L alEC/pmEC: grey matter density in left anterolateral/posteromedial entorhinal cortex voxel. AUC: area under the ROC curve. OT: optimal threshold. Sensitivity is percentage of correctly identified converters. Specificity is percentage of correctly identified controls. Accuracy is the percentage of all correct classifications. These values are provided for 3 different thresholds:

**CV'd**: Based on cross-validated thresholds (described in the Methods section of the main manuscript). The test data is split in half (21 converters and 21 controls), and the optimal cut-off threshold is decided based on the labels and ROC of these subjects. This threshold is then applied to remaining subjects to compute specificity and sensitivity, and this process then repeated for the other half. For the threshold, we report the median values of 2 x 50 split-half experiments. Performance measures (accuracy, sensitivity, specificity) are averages over these split-half experiments (hence the discrepancy between CV'd and OT performance measures despite the same threshold).

**OT**: Based on optimal thresholds. The labels in the test set are used to decide which cut-off threshold (converters *vs*. non-converters) is optimal in terms of (balanced) accuracy along the ROC curve (*i.e.*, which point of ROC curve gives the best accuracy). Since the labels in the test set are being used, these thresholds are decided based on the test data. While AUC is a proper test set-based measure, sensitivity and specificity describe how they would be if the logistic regression model was perfectly calibrated.

**0.5**: Setting the thresholds to 0.5. This is straight-forward application of the model, but produces poor sensitivity and specificity values as the model is not correctly calibrated for learning effects (*i.e.*, the effects of repeated yearly testing on neuropsychological test performance).

| | | | | |
|---|---|---|---|---|
| **Demographics**<br>Constant : 0.061<br>Age : -0.169<br>Gender : 0.220<br>Best Cross-validated AUC: 0.589;<br>Alpha:0.2; Lambda:0.13<br>Test AUC: 0.491<br>+ 95% CI [0.366, 0.615] | Threshold | **CVd** 0.516 | **OT** 0.492 | **0.5** |
| | Test sensitivity | 0.415 | 0.643 | 0.571 |
| | Test specificity | 0.499 | 0.405 | 0.429 |
| | Test accuracy | 0.457 | 0.524 | 0.500 |
| **Demographics plus APOE genotype**<br>Constant : 0.182<br>Age : -0.302<br>Gender : 0.327<br>APOE ε4 : -0.352<br>Education : 0.163<br>Best Cross-validated AUC: 0.641;<br>Alpha:0; Lambda:0.04<br>Test AUC: 0.647<br>+ 95% CI [0.515, 0.753] | Threshold | **CVd** 0.495 | **OT** 0.487 | **0.5** |
| | Test sensitivity | 0.572 | 0.738 | 0.619 |
| | Test specificity | 0.575 | 0.524 | 0.595 |
| | Test accuracy | 0.574 | 0.631 | 0.607 |
| **MRI**<br>Constant : 0.251<br>GMD L alEC (-26, -14, -32) : 0.0327<br>GMD L alEC (-26, -16, -30) : 0.0855<br>GMD L pmEC (-26, -18, -28) : 0.246<br>GMD L pmEC (-28, -20, -26) : 0.149<br>Best Cross-validated AUC: 0.825;<br>Alpha:0.86; Lambda:0.2<br>Test AUC: 0.800<br>+ 95% CI [0.684, 0.880] | Threshold | **CVd** 0.492 | **OT** 0.492 | **0.5** |
| | Test sensitivity | 0.716 | 0.738 | 0.643 |
| | Test specificity | 0.727 | 0.786 | 0.810 |
| | Test accuracy | 0.722 | 0.762 | 0.726 |
| **Neuropsychology**<br>Constant : 0.363<br>FAQ : -0.110<br>FCSRTrecdiftot : 0.460<br>Best Cross-validated AUC: 0.870;<br>Alpha:0.58; Lambda:0.19<br>Test AUC: 0.876<br>+ 95% CI [0.783, 0.938] | Threshold | **CVd** 0.399 | **OT** 0.399 | **0.5** |
| | Test sensitivity | 0.716 | 0.810 | 0.452 |
| | Test specificity | 0.825 | 0.810 | 0.952 |
| | Test accuracy | 0.771 | 0.810 | 0.702 |
| **Neuropsychology plus Demographics**<br>Constant : 0.391<br>FAQ : -0.118<br>FCSRTrecdiftot : 0.497<br>Best Cross-validated AUC: 0.870;<br>Alpha:0.63; Lambda:0.17<br>Test AUC: 0.876<br>+ 95% CI [0.782, 0.936] | Threshold | **CVd** 0.391 | **OT** 0.399 | **0.5** |
| | Test sensitivity | 0.716 | 0.810 | 0.452 |
| | Test specificity | 0.825 | 0.810 | 0.952 |
| | Test accuracy | 0.771 | 0.810 | 0.702 |
| **All Modalities**<br>Constant : 0.517<br>FCSRTrecdiftot : 0.397<br>GMD L pmEC (-26, -18, -28) : 0.304<br>GMD L pmEC (-28, -20, -26) : 0.185<br>Best Cross-validated AUC: 0.919;<br>Alpha:0.94; Lambda:0.16<br>Test AUC: 0.905<br>+ 95% CI [0.825, 0.954] | Threshold | **CVd** 0.428 | **OT** 0.428 | **0.5** |
| | Test sensitivity | 0.739 | 0.762 | 0.476 |
| | Test specificity | 0.880 | 0.929 | 0.976 |
| | Test accuracy | 0.809 | 0.845 | 0.726 |

**Supplementary Table 9. Statistical comparison of 5 statistical models on test data.** Upper Triangle: AUC differences. Bottom Triangle: *P*-values of AUC differences based on DeLong's test. In bold, *P* <0.05.

| | **All Modalities** | **Neuropsych -ology** | **MRI** | **Demographics plus APOE genotype** | **Demographics** |
|---|---|---|---|---|---|
| **All Modalities** | | 0.02891 | 0.10431 | 0.25737 | 0.39626 |
| **Neuropsychology** | 0.47338 | | 0.07540 | 0.22846 | 0.36735 |
| **MRI** | **0.00774** | 0.26259 | | 0.15306 | 0.29195 |
| **Demographics plus APOE genotype** | **0.00003** | **0.00077** | **0.03816** | | 0.13889 |
| **Demographics** | **0.00000** | **0.00000** | **0.00089** | 0.22334 | |

**Supplementary equation (1) and (2)**

All Modalities model with raw (unstandardized) variables:

$$\log\left(\frac{P(MCI)}{1-P(MCI)}\right) = 7.61 - 0.229 \times FCSRT - 5.15 \times MRI_{(-26,-18,-28)} - 0.1056 \times MRI_{(-28,-20,-26)} \quad (1)$$

Neuropsychology model with raw (unstandardized) variables:

$$\log\left(\frac{P(MCI)}{1-P(MCI)}\right) = 3.43 - 0.266 \times FCSRT - 0.146\, FAQ \quad (2)$$

The obtained value is then transformed into predicted probability using exp(*x*)/(1+exp(*x*)) transformation followed by assigning cases of predicted MCI development in 1 year for probabilities >0.5 and controls for probabilities <0.5.

**Supplementary Figure 1. Subdivisions of human EC show similar grey matter density reductions the year before MCI development. a.** Probabilistic maps of anterolateral and posteromedial entorhinal cortex have been warped onto each subjects' MRI, here shown for an example control subject in cyan and magenta, respectively, and the average GMD extracted. **b.** GMD values are plotted for Converters, Matched and Unmatched controls, with EC subdivision indicated by coloured bar above each plot. There is a significant main effect of group between Converters *vs*. Matched or Unmatched controls (**Supplementary Table 3)**. The central mark indicates the median, and the bottom and top box edges indicate the 25th and 75th percentiles, respectively. Whiskers extend 1.5 times the interquartile range away from the top or bottom of the box, and outliers (+) are plotted individually.

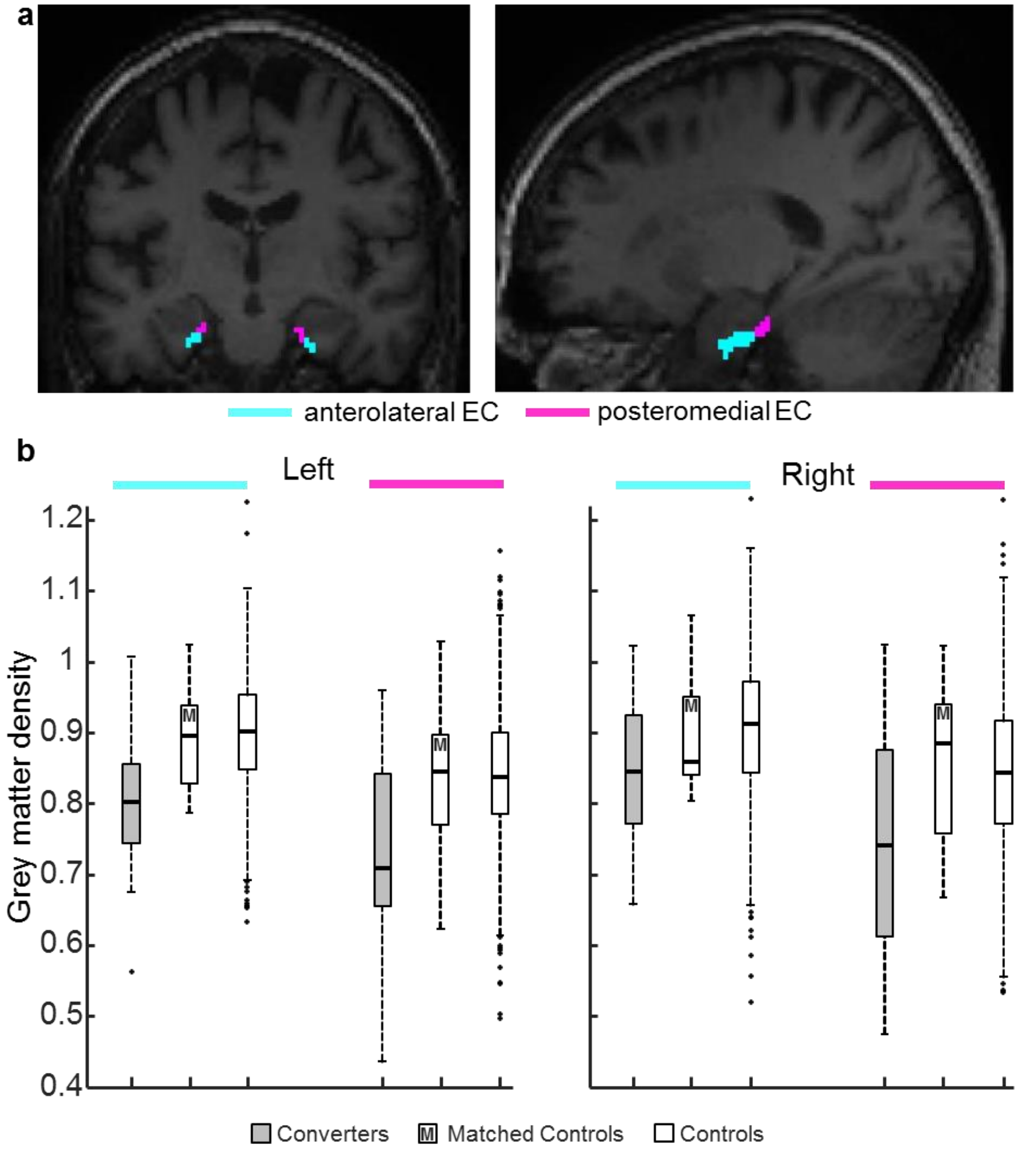

**Supplementary Figure 2**. **Medial temporal atrophy in healthy elderly individuals destined for MCI – test group.** Decreased grey matter density (GMD) at $V_{conv-1}$ for 42 validation converters *vs.* 776 controls is limited primarily to the medial temporal lobe. (**a**) Reduced GMD, selective to bilateral amygdala, hippocampus, and EC in converters one year before MCI onset, is depicted on serial coronal sections, overlaid on the group averaged T1 scan (threshold $P < 0.05$ whole-brain family-wise error corrected). (**b**) GMD in the global peak voxel (left entorhinal cortex; -26, -24, -22) is plotted for converters and controls. The central mark indicates the median, and the bottom and top box edges indicate the 25th and 75th percentiles, respectively. Whiskers extend 1.5 times the interquartile range away from the top or bottom of the box, and outliers (+) are plotted individually.

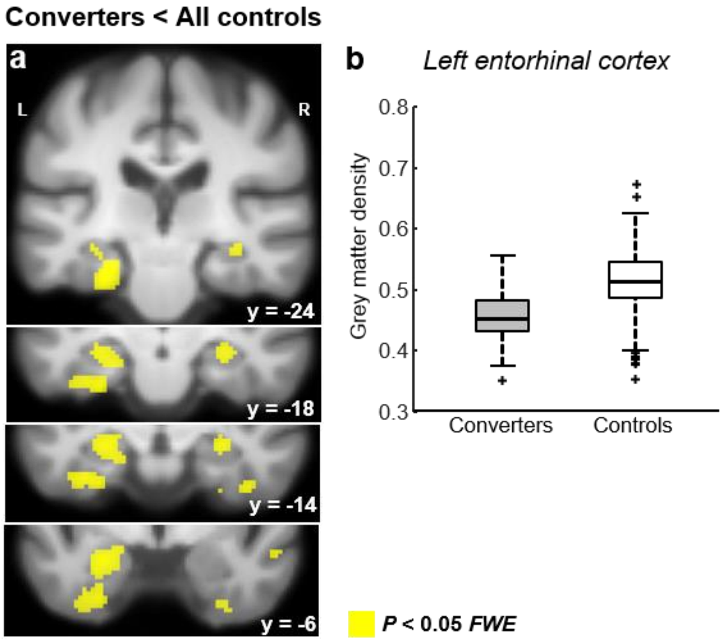
